# Overly strong priors for socially meaningful visual signals in psychosis proneness

**DOI:** 10.1101/473421

**Authors:** Heiner Stuke, Elisabeth Kress, Veith Andreas Weilnhammer, Philipp Sterzer, Katharina Schmack

## Abstract

Predictive coding accounts of psychosis state that an overweighing of high-level priors relative to sensory information may lead to the misperception of meaningful signals underlying the experience of auditory hallucinations and delusions. However, it is currently unclear whether the hypothesized overweighing of priors (1) represents a pervasive alteration that also affects the visual modality, and, (2) takes already effect at early automatic processing stages.

Here, we addressed these questions by studying visual perception of socially meaningful stimuli in healthy individuals with varying degrees of psychosis proneness (n=39). In a first task, we quantified participants’ prior for detecting faces in visual noise. In a second task, we measured participants’ prior for detecting direct gaze stimuli that were rendered invisible by continuous flash suppression. We found that the prior for detecting faces in noise correlated with hallucination proneness (rho=0.50, p=0.001) as well as delusion proneness (rho=0.44, p=0.005). Similarly, the prior for detecting invisible direct gaze was significantly associated with hallucination proneness (rho = 0.42, p = 0.010) and trend-wise with delusion proneness (rho = 0.29, p = 0.087). Our results provide evidence for the idea that overly strong high-level priors for automatically detecting socially meaningful stimuli might constitute a generic processing alteration in psychosis.

## INTRODUCTION

Schizophrenia is characterized by psychotic symptoms such as delusions and hallucinations. Neurocognitive theories that draw on predictive coding and Bayesian theories of brain function have proposed an imbalance between prior expectations and current sensory information as a central disturbance underlying psychotic experiences (Fletcher and Frith, 2009; Adams et al., 2013; Sterzer et al., 2018). In this context, an overly strong prior for socially meaningful signals can account for hallucinatory experiences such as hearing voices in the absence of causative stimulus, or delusional experiences such as the feeling of being looked at by strangers (Corlett et al., 2009).

Consistent with this theoretical framework, an increased tendency to perceive voices in auditory noise has been observed in psychosis and related conditions (Bentall and Slade, 1985; Hoffman et al., 2007; Vercammen et al., 2008; Galdos et al., 2011; Alderson-Day et al., 2017), in line with the idea of overly strong prior for socially meaningful signals in the auditory domain. A similar shift towards perceiving abstract signals such as pure tones in auditory noise (Powers et al., 2017), pointing to the possibility that overly strong priors might affect auditory perception in general.

Hence, whilst there is evidence to support the idea of overly strong priors for meaningful auditory signals in psychosis, it is currently unclear whether this reflects a generic processing deficits that reliably extends to the visual modality. A few studies have related an increased tendency to perceive faces in visual noise (Partos et al., 2016), and an increased tendency to perceive visual gaze as direct (Rosse et al., 1994; Hooker and Park, 2005; Tso et al., 2012) to psychosis and related conditions, but results have been mixed (see (Franck et al., 2002) for a negative report). Assessing relationships between psychotic experiences and the use of priors towards meaningful visual signals is crucial for probing the generalizability of strong prior accounts of psychosis. Here, we therefore related psychosis proneness in individuals from the general population to behavior in a visual detection-in-noise task. We hypothesized that psychosis proneness would positively correlate with the tendency to detect faces in visual noise, and hence a prior towards detecting meaningful stimuli.

Moreover, it is currently unclear which stage of information processing is affected by overly strong priors underlying psychotic experiences. It is conceivable that overly strong priors might only affect the late, conscious processing stage of cognitive interpretation. Alternatively, the effects of overly strong priors might extend to early, automatic sensory processing stages that determine the access of stimuli to awareness. In the visual domain, the potency of visual stimuli to gain access to awareness can be assessed with interocular masking techniques, such as continuous flash suppression (CFS) (Tsuchiya and Koch, 2005). In CFS, one eye is presented with a target stimulus, while the other eye is presented with a dynamic mask that initially suppresses the target stimulus from conscious perception. The time that the suppressed stimulus takes to to overcome interocular suppression has been proposed as a measure for the potency of a specific stimulus to gain access to awareness (Jiang et al., 2007; Stein and Sterzer, 2014). For example, this ‘breaking CFS’ paradigm (b-CFS; (Stein et al., 2011a) has been used to show that suppression times are decreased for stimuli with direct gaze as compared to stimuli with averted gaze (Stein et al., 2011b). Inter-individual variability in breakthrough time depends on individual factors related to the stimuli that compete for perceptual dominance. For example, the advantage for faces with direct gaze in gaining access to awareness is reduced in individuals with autistic traits (Madipakkam et al., 2018). Similarly, suppression times are reduced for sad faces in patients with major depression (Sterzer et al., 2011) and for spider stimuli in individuals with spider phobia (Schmack et al., 2016).

Here we asked whether a strong prior for direct gaze may affect those processing stages that determine access of face stimuli to awareness and therefore tested whether suppression times for direct compared to averted gaze may be shorter in individuals with high psychosis proneness.

## MATERIAL AND METHODS

### Participants and psychometry

Thirty-nine participants were recruited from the general population through advertising. The study was approved by the Ethical Committee of the Charité, Universitätsmedizin Berlin. After complete description of the study to the participants, written informed consent was obtained in accordance with the Declaration of Helsinki of 1975 before participation.

The participants' proneness to delusional ideation was quantified using the Peters Delusion Inventory, 21-item version (PDI-21, (Peters et al., 2004)). The 21 items of this self-rating questionnaire cover a wide range of delusional convictions, including beliefs in the paranormal, grandiosity ideas or suspicious thoughts. For every endorsed belief, the questionnaire asks for dimensional ratings of belief-related distress, preoccupation and conviction.

Additionally, proneness to hallucinatory experiences was assessed with the Cardiff anomalous perception scale (CAPS,(Bell et al., 2006). This 32-item self-rating scale assesses anomalous perceptual experiences in different sensory domains including proprioception, time perception, somatosensory perception, as well as visual and auditory perception. The intensity of every anomalous perception is quantified on subscales for intrusiveness, frequency and distress. As in our previous work (Stuke et al., 2017, 2018), we used total PDI and CAPS scores obtained by adding up their three subscales, and Spearman correlations to relate psychosis proneness to task behavior. Basic demographic information and average psychosis proneness scores of the participants are summarized in Table 1.

**Table 1.** Participants’ characteristics

| Characteristic | Mean (SD) |
| --- | --- |
| Age | 30.31 (10.06) |
| PDI score | 76.95 (42.86) |
| CAPS score | 106.90 (48.81) |
| Characteristic | Absolute numbers |
| Sex | female: 19; male: 20 |
| Smoking | yes: 11; no: 26; missing information: 2 |
| Vocation | None: 8; Apprenticeship: 1; Bachelor: 16; Master: 14 |

### Face task

To quantify priors towards socially meaningful stimuli in visual perception, we measured psychosis-like mispercepts of illusory faces in noise. To this end, we devised a face detection task that required the participants to detect faces embedded in noise. One-hundred stimuli (40 target and 60 noise stimuli) were created. Participants were instructed that a sequence of noisy stimuli will be presented to them and that some of stimuli will contain a human face. Each stimulus was presented for 3000 ms followed by a forced-choice decision of whether a face was present or not. After a response had been made and a subsequent inter trial interval of 800 ms, the next stimulus was presented (Figure 1).

**Figure 1.**
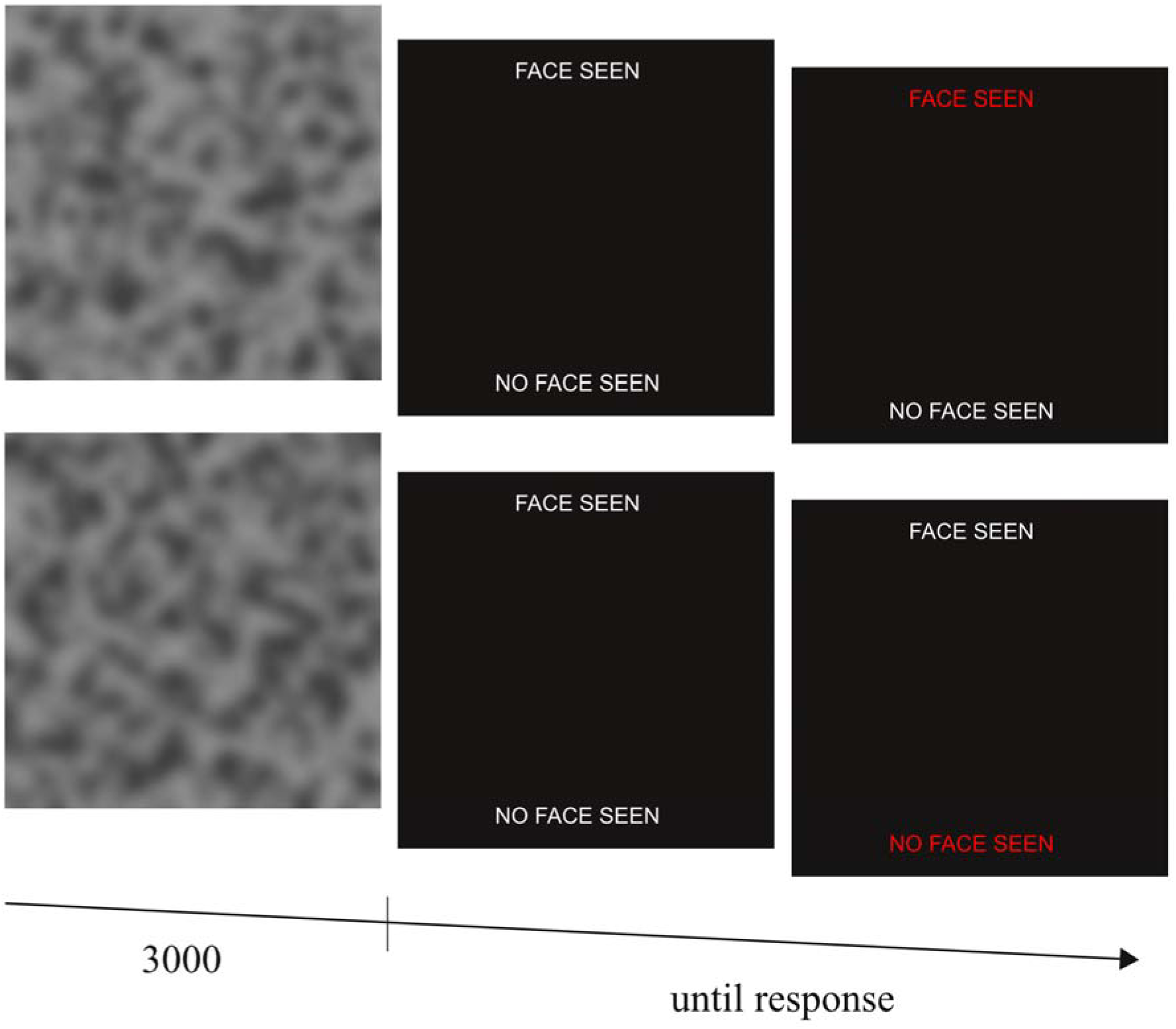
Experimental sequence of the face task for two exemplary trials with (top row) and without (lower row) embedded face. Face images were shown for 3000 ms followed by a binary forced choice indication of whether a face had been detected by the participants.

Stimuli were designed to resemble those that have proven to induce psychosis-like percepts of illusory faces in previous work (Partos et al., 2016). Noise stimuli consisted of a noise pattern only (without embedded face) and were created in three steps using Matlab and the Image Processing Toolbox. Firstly, basic noise patterns were generated by randomly placing a total of 1000 black circles with diameters varying randomly from 1 to 15 pixels (0.04° to 0.64° of visual angle) on a white image of 450 * 450 pixels (19.45° of visual angle). Secondly, the basic noise patterns were degraded by adding multiplicative noise (as implemented in the 'speckle' command of the Matlab imnoise routine with a distribution variance of 2). Finally, the resulting noise stimuli were blurred with a Gaussian filter ('gaussian' command of the Matlab imnoise routine with a distribution variance of 10) and image contrast was reduced with the 'imadjust' routine (resetting greyscale intensities to values between 0.1 and 0.9). For the target stimuli, twenty adult faces with neutral expression were taken from the Productive Ageing Laboratory Face Database (Minear and Park, 2004) and placed at random positions in the noise stimuli before the third step of noise image generation (i.e., before the Gaussian filter and contrast reduction). The specific image generation parameters were chosen to ensure that participants were imperfectly able to distinguish the faces from the noise stimuli in a pilot study with five participants (discriminability mean = 0.81, SD = 0.03; bias = 1.52, SD = 0.86; see below for details on computation of these measures).

### Face task analysis

Face task behavior was assessed with signal detection theory, which distinguishes a *bias* towards positive responses (i.e., to detect faces) from a *discriminability* measure d’ (i.e., a general ability to differentiate between faces and noise). The discriminability d’ is given by:

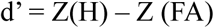

where H is the hit rate (percentage of stimuli with face, where a face was correctly detected), FA is the false alarms rate (percentage of stimuli without face, where a face was wrongly detected), and Z is the inverse of the cumulative distribution function of the Gaussian distribution. The corresponding bias measure was given by

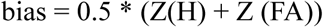

A high bias value reflects a tendency to detect faces in both noise and target stimuli, and hence an enhanced prior for meaningful information in the visual domain.

### Gaze task

To quantify the effect of individual priors for socially meaningful information on the access of visual stimuli to awareness, we used an established interocular suppression task with face stimuli that displayed either direct or averted gaze (Stein et al., 2011b; Seymour et al., 2016){Seymour, 2016 #1;Stein, 2011 #3}(Madipakkam et al., 2018). In this task, stimuli were photographs of three different female faces, each in a version with direct and averted gaze. The impression of eye gaze being either directed at or away from the observer was achieved by a shift of the pupil to the left or the right. For example, a head rotated to the right together with the pupil shifted to the left resulted in the impression of a face looking at the observer (see Fig. 2, lower left). All faces were cut into oval shapes comprising a size of 3.8° × 4.5° and equalized for global contrast and luminance. Participants viewed the screen through a mirror stereoscope, which provided separate visual input to the two eyes. The participant’s head was stabilized by a chin rest at a viewing distance of 50 cm and stimuli were displayed on a 19- in. CRT monitor (resolution: 1024 × 768 Px; refresh rate: 60 Hz).

**Figure 2.**
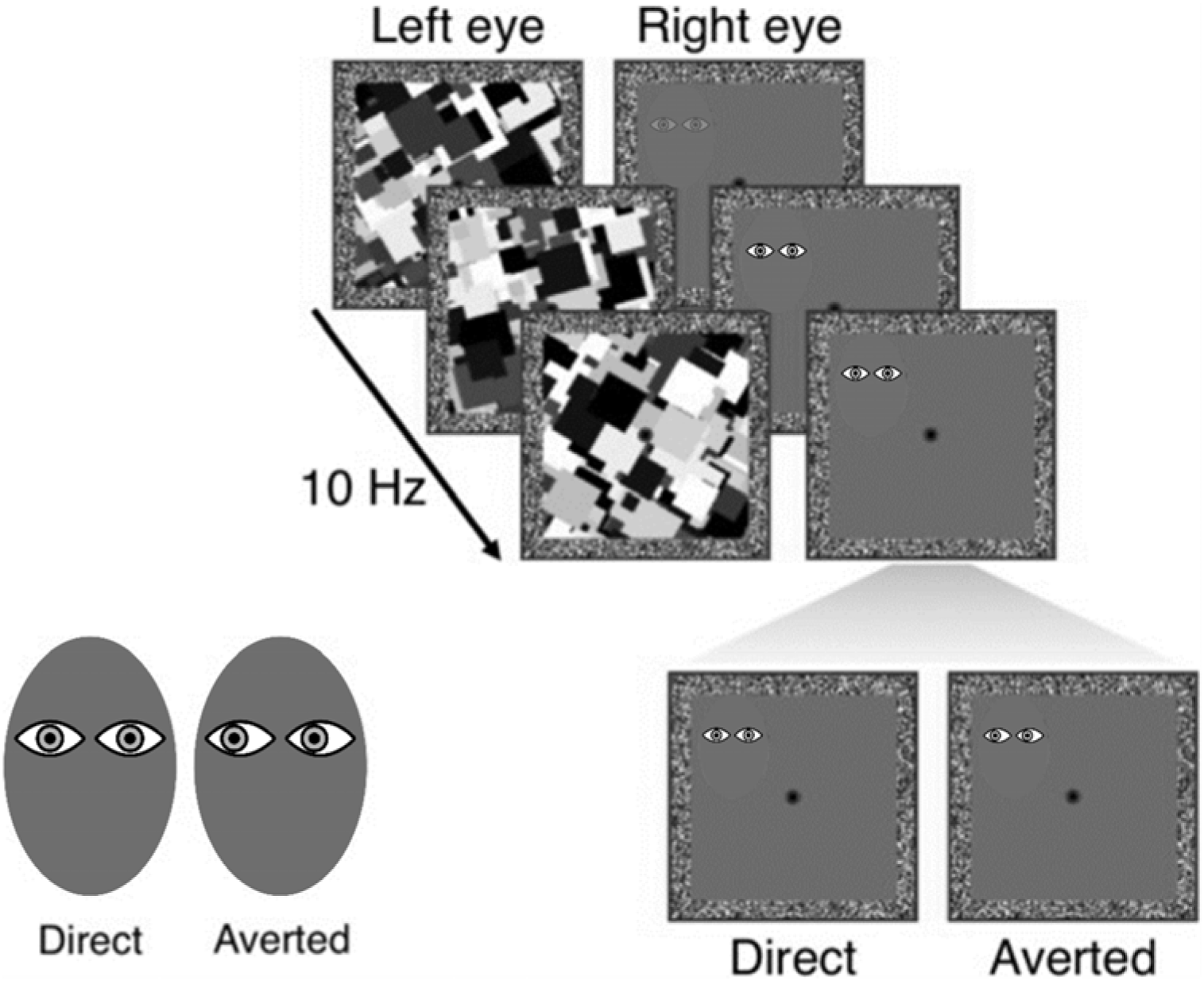
Experimental sequence of the unconscious gaze task. Both eyes of the participants received separate stimulation through a mirror stereoscope. One eye was shown the target stimulus (photograph of faces with either direct or averted gaze, here shown in symbolized version), whose conscious perception was suppressed by a dynamical mask shown to the other eye. Participants were instructed to indicate the localization of a face as soon as it broke through the mask (but not told that faces differed with respect to gaze). By subtracting mean response times for averted gaze from mean response times for direct gaze, we obtained a measure of the priority of unconscious processing of direct gaze.

The effect of eye gaze on access of face stimuli to awareness was assessed using bCFS. Each trial began with a 2 s presentation of white frames (12.0° × 12.0°) with a grey background and a red fixation cross (Figure 2). Thereafter, high-contrast, grey scale, dynamic masks were flashed to a randomly selected eye at a frequency of 10Hz while simultaneously a face stimulus with either a direct or averted gaze was gradually introduced to the other eye. The contrast of the face stimulus was gradually increased from 0% to 100% within the first second from the beginning of the trial and the stimulus remained at maximum contrast until a response was made or for a maximum of 15 s. The stimuli could be presented in one of the four quadrants of the white frame (3.4° horizontal displacement from the fixation cross and 3° vertical displacement). Participants had to indicate the location of the face (i.e., the quadrant) by button press as soon as the face overcame interocular suppression. Importantly, participants’ task (i.e. location discrimination) was orthogonal to the condition of interest (i.e. gaze direction of the presented faces). Participants were therefore unaware of the existence of two different gaze directions. The eye to which the face stimulus and masks were presented were randomized and counter-balanced. The eyes viewing the masks and the face stimuli, respectively, were randomized across trials. Participants completed a total of 48 trials, 24 for with direct and 24 with averted gaze. Target variables were the response (breakthrough) times for correctly localized faces.

### Gaze task analysis

Three of the 39 participants were not included in the gaze task analysis, one because of technical problems and two because the task did not work due to excessive stimulus suppression by the mask (more than 65% missed trials).

Analogously to previous studies using the same task, we compared mean breakthrough times separately for faces with direct and averted gaze. By subtracting breakthrough times for direct gaze from breakthrough times for averted gaze, we obtained a measure of the tendency towards faster access to awareness of direct gaze. In the following, we denote this measure as ‘direct gaze bias’, where a positive value indicates shorter breakthrough times for direct gaze relatively relative to averted gaze. As a sanity check, we first tested if this measure was significantly above zero (one sample t test), e.g., whether we could replicate previous findings of a generally faster breakthrough of direct gaze. Secondly, we tested whether the degree of this direct gaze bias depended on the individual’s psychosis proneness by correlating it with CAPS and PDI scores.

## RESULTS

In the face task, the bias for face detection correlated with both hallucination proneness (rho = 0.500, p = 0.001, n = 39) and delusion proneness (rho = 0.444, p = 0.005, n = 39). These results suggest that psychosis proneness is associated with an increased prior for faces in a detection-in-noise task (Figure 3a). In contrast, the discriminability measure d’ was not significantly related to hallucination proneness (rho = −0.087, p = 0.599, n = 39) or delusion proneness (rho = −0.184, p = 0.261, n = 39). Thus, there was no evidence for a significant association of psychosis proneness with the ability to discriminate between face and noise stimuli.

**Figure 3:**
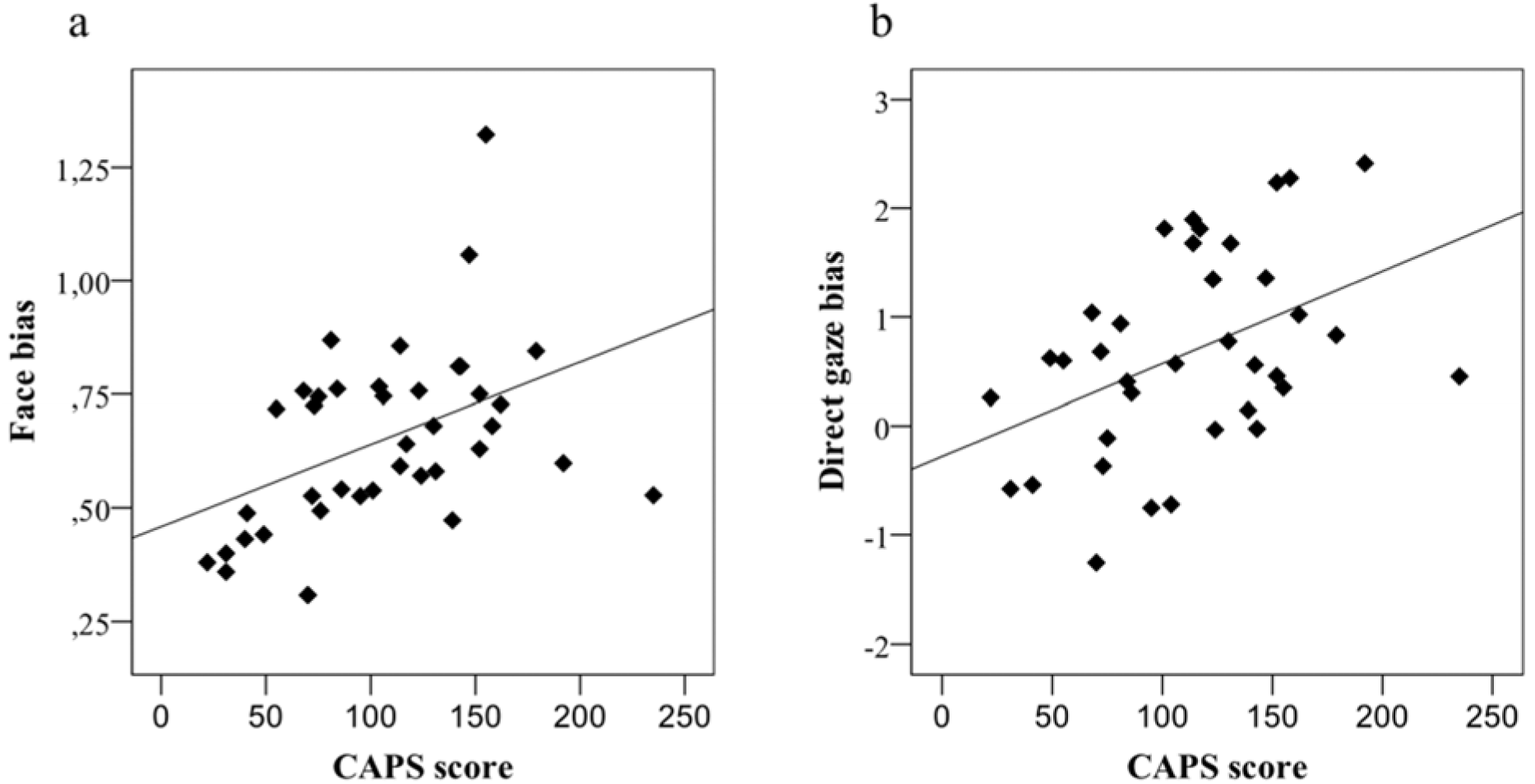
Relationships between participants’ hallucination proneness (CAPS scores) and signal detection bias in the face task (a) and direct gaze bias in the unconscious gaze task (b). With growing hallucination proneness, participants show an increased readiness to detect faces in noise and actual face stimuli as well as to unconsciously process direct gaze faster than averted gaze.

In the gaze task, breakthrough times were overall significantly shorter for direct compared to averted gaze (paired t test, T = −4.362, p < 0.001). Consistent with previous work (Stein et al., 2011b; Seymour et al., 2016), this result indicates a general direct-gaze bias for access to awareness in the whole sample (Figure 4). Importantly, this direct gaze bias correlated significantly with hallucination proneness (CAPS scores, Spearman rank correlation, rho = 0.424, p = 0.010, n = 36) and trend-wise with delusion proneness (PDI scores, rho = 0.289, p = 0.087, n = 36). These results show that psychosis proneness is associated with enhanced access of direct gaze to awareness, suggesting a stronger a prior for socially relevant visual information (Figure 3b).

**Figure 4:**
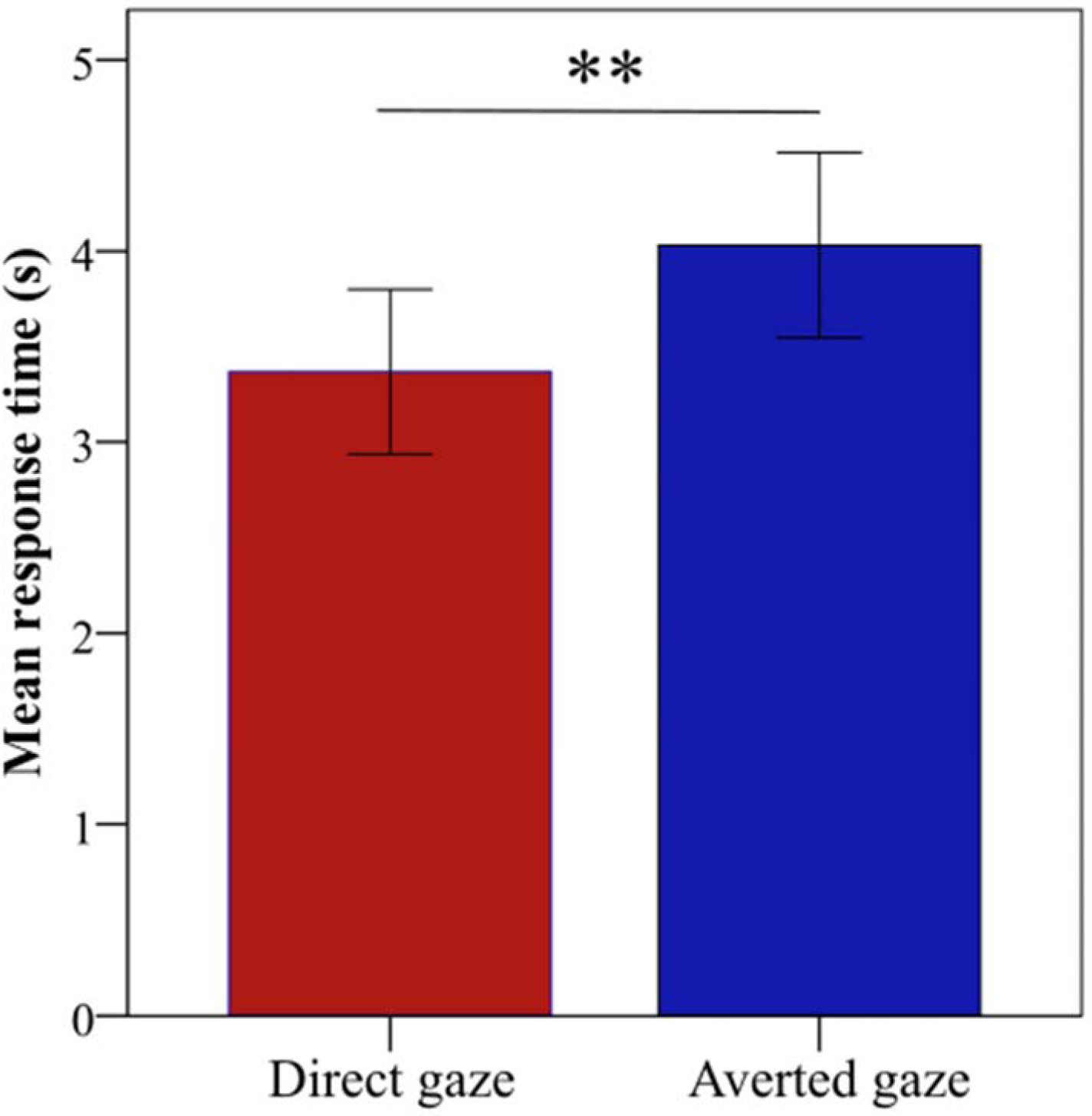
Mean response times for direct and averted gaze in the unconscious gaze task. Consistent with previous studies, response times were significantly faster for direct gaze (T = −4.362, p < 0.001).

## DISCUSSION

In the present work, we demonstrate an increased prior for socially meaningful information in noisy or perceptually masked visual stimuli as a correlate of psychosis proneness in healthy people. These results are compatible with strong prior accounts that state a “superimposed” prior on naturally noisy perception as a substrate for psychotic experiences (Corlett et al., 2009).

It should be noted that, contrary to other study designs, “prior” does not refer to an experimentally manipulated information in our experiments, but to an expectation of meaningful signal (faces, direct gaze) in noisy and ambiguous stimuli. In previous experiments in which prior information was varied and had to be balanced against (potentially contradictory) sensory information, relationships were less clear and some studies even reported *decreased* prior usage with growing psychotic experiences (Jardri et al., 2017; Stuke et al., 2018). This inconsistency goes well with the emerging understanding that different kinds of priors may be differentially affected in psychosis, and that alterations in perceptual inference go beyond a simple over- or underweighting of priors (for a detailed discussion, see (Sterzer et al., 2018).

Our finding of a relationship between hallucination proneness and an increased prior for direct gaze in a masking task is of relevance for the ongoing debate about the processing stage, at which psychosis-typical perceptual alterations take effect (Berkovitch et al., 2017). Here, our finding speaks for an involvement of early, automatic processing stages. However, there is disagreement with a negative finding of (Seymour et al., 2016), who assessed differences between schizophrenia patients and healthy controls in a very similar task and did not find any significant differences between the groups. One explanation for these conflicting findings might be that an enhanced unconscious direct gaze prior is present in people at-risk for psychosis, but not in the chronic, stable and medicated sample investigated by Seymour et al., 2016. Another reason might lie in slightly different properties of the CFS tasks used in this study and in the study by Seymour et al.. For instance, in the task used by Seymour et al., the mask intensity was gradually decreased after target stimulus fade-in, whereas in our task version, the mask intensity did not change. Gradual fade-out of the mask may render bCFS less sensitive for the detection of individual differences (Münkler et al., 2015). Nevertheless, the mean breakthrough times as well as gaze-dependent breakthrough time differences lie in a similar range in both studies, rendering major differences in task-elicited effects unlikely. Finally, it should be noted that Seymour et al. also report increased response time differences in patients, which however did not reach significance.

False alarms in detection tasks (perception of meaning from noise stimuli) have an intuitive “face validity” as experimental hallucination markers. Similarly, the preferential unconscious processing of direct gaze directly relates to the delusional feeling of being stared at in public. Hence, as opposed to other common markers for psychosis proneness (e.g., a reduced EEG mismatch negativity (Näätänen et al., 2015; Erickson et al., 2016) and cognitive biases such as jumping-to-conclusions (Dudley et al., 2016), the two tasks used here have an immediate connection to the phenomenology of psychosis and might serve as symptom-related markers for the severity of psychotic experiences. It might be a worthwhile endeavor to investigate the predictive power of these markers in further research. In clinics, an early response of psychosis-related markers after initiation of antipsychotic treatment might help to predict following treatment response (up to now, early treatment response remains the most reliable predictor of long-term treatment outcome (Correll et al., 2003; Kinon et al., 2008; Stauffer et al., 2011). In animal research, similar detection-in-noise tasks might help to assess the effects of pro- or antipsychotic interventions. In any case, the development of suited experimental markers to monitor and predict the effect of psychosis-targeting interventions remains an important cornerstone for progressing our still limited understanding and treatment options for psychotic disorders.

In summary, our results speak to an overly strong prior for socially meaningful information in people with psychotic experiences that extends beyond the domain of auditory perception and likely affects early unconscious stages of sensory processing.

## REFERENCES

Adams RA, Stephan KE, Brown HR, Frith CD, Friston KJ (2013) The computational anatomy of psychosis. Front Psychiatry 4: 47.

Alderson-Day B, Lima CF, Evans S, Krishnan S, Shanmugalingam P, Fernyhough C, Scott SK (2017) Distinct processing of ambiguous speech in people with non-clinical auditory verbal hallucinations. Brain J Neurol 140:2475–2489.

Bell V, Halligan PW, Ellis HD (2006) The Cardiff Anomalous Perceptions Scale (CAPS): a new validated measure of anomalous perceptual experience. Schizophr Bull 32:366–377.

Bentall RP, Slade PD (1985) Reality testing and auditory hallucinations: a signal detection analysis. Br J Clin Psychol 24 (Pt 3):159–169.

Berkovitch L, Dehaene S, Gaillard R (2017) Disruption of Conscious Access in Schizophrenia. Trends Cogn Sci 21:878–892.

Corlett PR, Frith CD, Fletcher PC (2009) From drugs to deprivation: a Bayesian framework for understanding models of psychosis. Psychopharmacology (Berl) 206:515–530.

Correll CU, Malhotra AK, Kaushik S, McMeniman M, Kane JM (2003) Early prediction of antipsychotic response in schizophrenia. Am J Psychiatry 160:2063–2065.

Dudley R, Taylor P, Wickham S, Hutton P (2016) Psychosis, Delusions and the “Jumping to Conclusions” Reasoning Bias: A Systematic Review and Meta-analysis. Schizophr Bull 42:652–665.

Erickson MA, Ruffle A, Gold JM (2016) A Meta-Analysis of Mismatch Negativity in Schizophrenia: From Clinical Risk to Disease Specificity and Progression. Biol Psychiatry 79:980–987.

Fletcher PC, Frith CD (2009) Perceiving is believing: a Bayesian approach to explaining the positive symptoms of schizophrenia. Nat Rev Neurosci 10:48–58.

Franck N, Montoute T, Labruyère N, Tiberghien G, Marie-Cardine M, Daléry J, d’Amato T, Georgieff N (2002) Gaze direction determination in schizophrenia. Schizophr Res 56:225–234.

Galdos M, Simons C, Fernandez-Rivas A, Wichers M, Peralta C, Lataster T, Amer G, Myin-Germeys I, Allardyce J, Gonzalez-Torres MA, van Os J (2011) Affectively salient meaning in random noise: a task sensitive to psychosis liability. Schizophr Bull 37:1179–1186.

Hoffman RE, Woods SW, Hawkins KA, Pittman B, Tohen M, Preda A, Breier A, Glist J, Addington J, Perkins DO, McGlashan TH (2007) Extracting spurious messages from noise and risk of schizophrenia-spectrum disorders in a prodromal population. Br J Psychiatry J Ment Sci 191:355–356.

Hooker C, Park S (2005) You must be looking at me: the nature of gaze perception in schizophrenia patients. Cognit Neuropsychiatry 10:327–345.

Jardri R, Duverne S, Litvinova AS, Denève S (2017) Experimental evidence for circular inference in schizophrenia. Nat Commun 8:14218.

Jiang Y, Costello P, He S (2007) Processing of invisible stimuli: advantage of upright faces and recognizable words in overcoming interocular suppression. Psychol Sci 18:349–355.

Kinon BJ, Chen L, Ascher-Svanum H, Stauffer VL, Kollack-Walker S, Sniadecki JL, Kane JM (2008) Predicting response to atypical antipsychotics based on early response in the treatment of schizophrenia. Schizophr Res 102:230–240.

Madipakkam AR, Rothkirch M, Dziobek I, Sterzer P (2018) Access to awareness of direct gaze is related to autistic traits. Psychol Med:1–7.

Minear M, Park DC (2004) A lifespan database of adult facial stimuli. Behav Res Methods Instrum Comput J Psychon Soc Inc 36:630–633.

Münkler P, Rothkirch M, Dalati Y, Schmack K, Sterzer P (2015) Biased recognition of facial affect in patients with major depressive disorder reflects clinical state. PloS One 10:e0129863.

Näätänen R, Shiga T, Asano S, Yabe H (2015) Mismatch negativity (MMN) deficiency: a break-through biomarker in predicting psychosis onset. Int J Psychophysiol Off J Int Organ Psychophysiol 95:338–344.

Partos TR, Cropper SJ, Rawlings D (2016) You Don’t See What I See: Individual Differences in the Perception of Meaning from Visual Stimuli. PloS One 11: e0150615.

Peters E, Joseph S, Day S, Garety P (2004) Measuring delusional ideation: the 21-item Peters et al. Delusions Inventory (PDI). Schizophr Bull 30:1005–1022.

Powers AR, Mathys C, Corlett PR (2017) Pavlovian conditioning-induced hallucinations result from overweighting of perceptual priors. Science 357:596–600.

Rosse RB, Kendrick K, Wyatt RJ, Isaac A, Deutsch SI (1994) Gaze discrimination in patients with schizophrenia: preliminary report. Am J Psychiatry 151:919–921.

Schmack K, Burk J, Haynes J-D, Sterzer P (2016) Predicting Subjective Affective Salience from Cortical Responses to Invisible Object Stimuli. Cereb Cortex N Y N 1991 26:3453–3460.

Seymour K, Rhodes G, Stein T, Langdon R (2016) Intact unconscious processing of eye contact in schizophrenia. Schizophr Res Cogn 3:15–19.

Stauffer VL, Case M, Kinon BJ, Conley R, Ascher-Svanum H, Kollack-Walker S, Kane J, McEvoy J, Lieberman J (2011) Early response to antipsychotic therapy as a clinical marker of subsequent response in the treatment of patients with first-episode psychosis. Psychiatry Res 187:42–48.

Stein T, Hebart MN, Sterzer P (2011a) Breaking Continuous Flash Suppression: A New Measure of Unconscious Processing during Interocular Suppression? Front Hum Neurosci 5: 167.

Stein T, Senju A, Peelen MV, Sterzer P (2011b) Eye contact facilitates awareness of faces during interocular suppression. Cognition 119:307–311.

Stein T, Sterzer P (2014) Unconscious processing under interocular suppression: getting the right measure. Front Psychol 5: 387.

Sterzer P, Adams RA, Fletcher P, Frith C, Lawrie SM, Muckli L, Petrovic P, Uhlhaas P, Voss M, Corlett PR (2018) The Predictive Coding Account of Psychosis. Biol Psychiatry 84:634–643.

Sterzer P, Hilgenfeldt T, Freudenberg P, Bermpohl F, Adli M (2011) Access of emotional information to visual awareness in patients with major depressive disorder. Psychol Med 41:1615–1624.

Stuke H, Stuke H, Weilnhammer VA, Schmack K (2017) Psychotic Experiences and Overhasty Inferences Are Related to Maladaptive Learning. PLoS Comput Biol 13: e1005328.

Stuke H, Weilnhammer VA, Sterzer P, Schmack K (2018) Delusion Proneness is Linked to a Reduced Usage of Prior Beliefs in Perceptual Decisions. Schizophr Bull.

Tso IF, Mui ML, Taylor SF, Deldin PJ (2012) Eye-contact perception in schizophrenia: relationship with symptoms and socioemotional functioning. J Abnorm Psychol 121:616–627.

Tsuchiya N, Koch C (2005) Continuous flash suppression reduces negative afterimages. Nat Neurosci 8:1096–1101.

Vercammen A, de Haan EHF, Aleman A (2008) Hearing a voice in the noise: auditory hallucinations and speech perception. Psychol Med 38:1177–1184.

